# Flexible representation of higher-dimensional cognitive variables with grid cells

**DOI:** 10.1101/578641

**Authors:** Mirko Klukas, Marcus Lewis, Ila Fiete

**Affiliations:** MIT, Cambridge, MA; Numenta, Redwood City, CA

## Abstract

We shed light on the theoretical capabilities of entorhinal grid cells to encode variables of dimension greater than two. Our model constructs representations of high-dimensional inputs through a combination of low-dimensional random projections and “classical” low-dimensional hexagonal grid cell responses. Without reconfiguration of the recurrent circuit, the same system can flexibly encode multiple variables of different dimensions while maximizing the coding range (per dimension) by automatically trading-off dimension with an exponentially large coding range. In contrast to previously proposed schemes, the model does not require the formation of higher-dimensional grid responses, a cell-inefficient and rigid mechanism. The firing fields observed in flying bats or climbing rats can be generated by neurons that combine activity from multiple grid modules, each representing higher-dimensional spaces according to our model. The idea expands our understanding of grid cells, suggesting that they could implement a general circuit that generates on-demand coding and memory states for variables in high-dimensional vector spaces.

## 1 Introduction

It is widely believed that entorhinal grid cells in mammals play a central role in the representation of spatial information. But recent evidence indicates that grid cells are more versatile than initially assumed and also represent cognitive variables other than (self-)location in physical space. They respond to the location of visual gaze [16, 20, 15], the locus of covert attention [25], or the values of two parametrically varied features of cartoon bird images [6]. In all these cases, the recorded cells exhibit a response structure that matches that of grid cells during spatial exploration, with single unit recordings in rodents indicating that the same grid cells are reused across variable types [17, 1]. This suggests that all of these variable types are represented by a single population of grid cells, which underlie very general types of cognitive representation. All these examples involve two-dimensional (2D) variables. However, cognitive variables aren’t limited to 2D, and it is a natural question to consider what kinds and dimensions of variables, theoretically, it is possible for grid cells to represent.

At the same time grid cell responses are structurally and dynamically constrained. Across a range of novel, familiar, and distorted spatial environments [28], during navigation on sloped terrains [11, 12] or one-dimensional tracks [29], and most strikingly, across sleep states when the animal receives no external spatial inputs and is rather driven by presumably high-dimensional internal spontaneous activity [8, 23], grid cells are confined to a fixed set of states with preserved cell-cell correlations that match those measured during awake exploration in familiar spatial environments. The fact that across coding states including sleep, grid cells conserve the pairwise firing relationships they exhibited in their spatial responses directly shows that the dynamics of a grid module are confined to a 2-dimensional set of states that is invariant across time, task, and behavioral state. Even the physical layout of grid cells in the brain is organized in a grid-like topographical pattern [13, 10] that mirrors – and likely drives – the functional response of grid modules. But how generally useful can the grid code be, if the autonomous states of each grid module are inherently 2-dimensional?

We propose a coding scheme for high-dimensional variables that is consistent with these structural and dynamical constraints and assume that the activity of each grid module is confined to a 2-dimensional toroidal attractor in the associated neural state space. From a purely mathematical viewpoint, the possibility of encoding higher-dimensional variables using grid cells is not surprising – after all the combined state space of multiple (M) 2-dimensional toroidal attractor manifolds, formed from the 2D grid responses of the *M* individual modules, is already a high-dimensional (2*M*-dimensional) toroidal manifold. However, the current literature does not offer any concrete coding scheme that exploits this fact. On the contrary, existing proposals and experimental searches center around the formation of grids that are themselves high-dimensional [14, 9].

These models face two major problems. The first is a question of resources. The formation of high-dimensional modules is cell-inefficient. For instance, a conservative estimate for the number of cells needed to form a single stable 3-dimensional continuous attractor network with resolution *K* per dimension is *K*^3^ while the same state capacity can be achieved by 3*K* cells by forming three lower dimensional attractor networks representing one dimension each.

The second is a question of flexibility. The recurrent connectivity of an attractor network is tailored to the dimension and geometry of the attractor manifold and cannot be easily reconfigured on demand. This is particularly problematic if the actual dimension of the input might vary; what if the circuit encodes a variable that appears to be lowdimensional but eventually turns out to vary in more dimensions than initially expected, or if the circuit must represent variables of different dimensions at different times?

In an ideal setting, coding states should be recruited flexibly to enable a large representational range or to encode additional dimensions, depending on the current demand (without changing previously assigned code words). Our model achieves this by combining random projections with a modulo arithmetic-like code based on “classic” grid cells. It naturally trades off representational range in one dimension for the ability to represent another and it does this with fixed recurrent connections.

## 2 Results

### 2.1 The grid code in 2D

Mammalian grid cells are defined by their periodic firing fields in planar environments: they fire at multiple locations corresponding to the vertices of an equilateral triangular lattice (Fig. 1a). A grid module is a discrete sub-network of grid cells with common underlying lattices (same period and orientation) that differ only by translational shifts. The firing fields of all the cells within a module uniformly cover the entire space such that at any time some sub-population within the module is active. Because of the periodically repeating responses of all cells within the module, together they represent the animal’s time-varying location *x*(*t*) as a 2-dimensional phase *ϕ*(*x*). This phase lies on a 2-dimensional torus in the neural state space [7].

**Figure 1:**
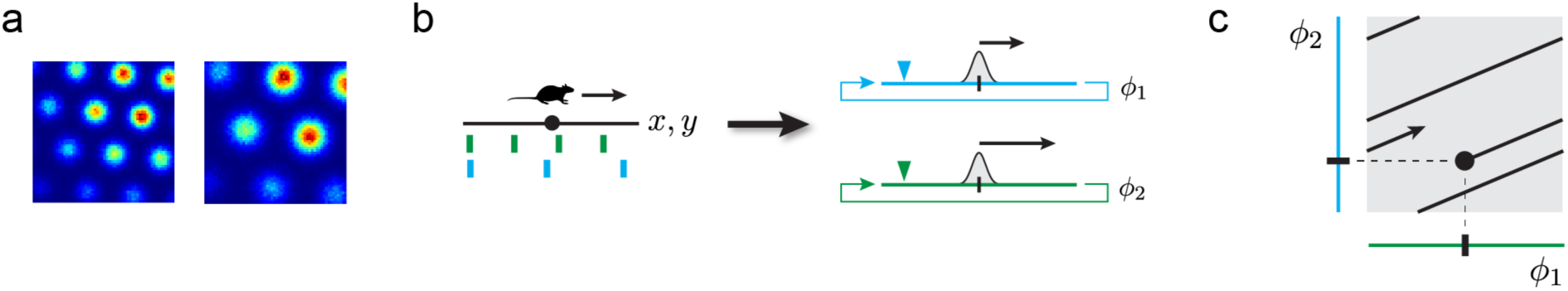
**A.** Firing fields and their hexagonal arrangement shown for two simulated grid cells from two modules of different scale. **B.** Schematic picture of the origin of periodic firing fields. Left: a schematic environment (black line) with cell responses (green, blue) of two grid cells from two modules of different scale. Right: schematic picture of two grid modules depicted as 1-dimensional circular continuous attractors. The colored triangles symbolize recording devices whose responses are shown on the left. A positional change (small arrow on left-hand side) corresponds to a change in phase (small arrows on right-hand side) in each module. Both phase-changes are related by a fixed scalar factor resulting in different spatial periodicity. **C.** Schematic picture of the coding space (gray box) spanned by multiple, here 2, grid modules (blue and green). The modulo-arithmetic nature of the grid code enables an extraordinarily huge representational range by tightly “packing/folding” an environment (black line) into code space (gray). The fixed ratio of module scales results in a linear embedding with fixed “slope”.

A single grid module, because of its periodic response, only provides an ambiguous estimate of the animal’s position. However decoding the position is possible if we consider the combined activity (or associated phases) of multiple modules whose lattices are distinct in spatial period. The range over which multiple modules resolve ambiguity is extremely large and reaches far beyond the scale of each individual lattice [7, 21]. We refer to the combined activity of multiple distinct modules, or equivalently the set of phases, as a grid code; Figure 1 provides a schematic picture illustrating the key mechanism and properties of the “classic” grid code. Before we go into more detail about capacity and representational range of the grid code we remind the reader how module activity is updated. From there defining our model is straightforward.

### 2.2 Velocity integration for instantaneous representation of arbitrary 2D variables

Central to key (attractor) models of grid cell dynamics [3], grid cell activity is updated based on estimated changes in animal position. Specifically, the phase (*ϕ*) encoded by a module is changed in proportion to the animal’s estimated instantaneous velocity (*v*):

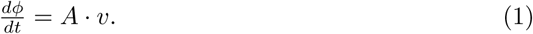

Here *A* ϵ ℝ^2*×*2^ is a linear operator that governs how the animal’s movements in the planar environment correspond to changes of the module’s internal phase.

Once the module is anchored, by assignment of a particular phase *φ*_0_ to a specific position *x*_0_ in the environment (e.g. through place cells or other spatially-specific cells [24, 4]), these velocity-based updates will automatically generate a grid code (phase) for any other location *x* in the environment reached via a path *γ* connecting *x* and *x*_0_:

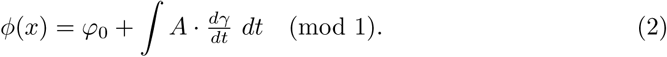

If the locations *x* lie in Euclidean space, the assigned phase is guaranteed to be independent of the particular path *γ* taken between *x*_0_ and *x*, thus ensuring a well-defined grid code for *x* regardless of trajectory:

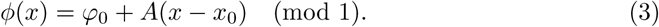

In this formulation, the different periods across modules could be generated by simple re-scaling of the velocity projection *A* by a gain factor. In fact, within a single module, when the spatial response period rapidly re-scales due to environment effects [2], the attractor model predicts that the re-scaling *must* be generated by a gain change in the velocity projection [3], rather than by a change in the recurrent wiring that gives rise of the periodic grid pattern. This prediction was verified in [28].

The description given above for spatial representation by grid cells, applies immediately to the local representation of arbitrary (locally Euclidean) 2D spaces: the grid phase is anchored to a particular landmark or landmarks in that space, and phase updates are made from motion cues generated by navigation through those spaces, automatically generating self-consistent codewords for any new location in that space. The only required change to represent a new space is the construction of a separate feedforward projection *A* for motion through that space; under this expanded view, existing grid cell models explain non-spatial representations [6, 16, 20, 15, 25, 17, 1].

### 2.3 Extension to arbitrary dimension: Our model

If grid cell activity in a module is inherently two-dimensional and modules cannot individually generate higher-dimensional grid responses, one way for the system to generate a response while navigating through higher-dimensional cognitive spaces is to simply project the *N*-dimensional velocity to two dimensions. We accomplish this by replacing the 2 × 2 matrix *A* in Equation 2 by one of shape 2 × *N*. Intuitively one can imagine the grid cells in a module responding to the variable’s “shadow” on (or projection onto) some plane in the cognitive space, and generating classical hexagonal grids (along that plane). Away from the plane the responses are elongated and undifferentiated along the directions of projecting (Fig. 2a,b,d). This scheme is consistent with the empirical findings in [11]. The resulting representation within a single module is ambiguous – at least *N* − 2 cognitive directions project to zero under *A* – and the cognitive variable is thus not decodable over any range, including within the smallest module period.

**Figure 2:**
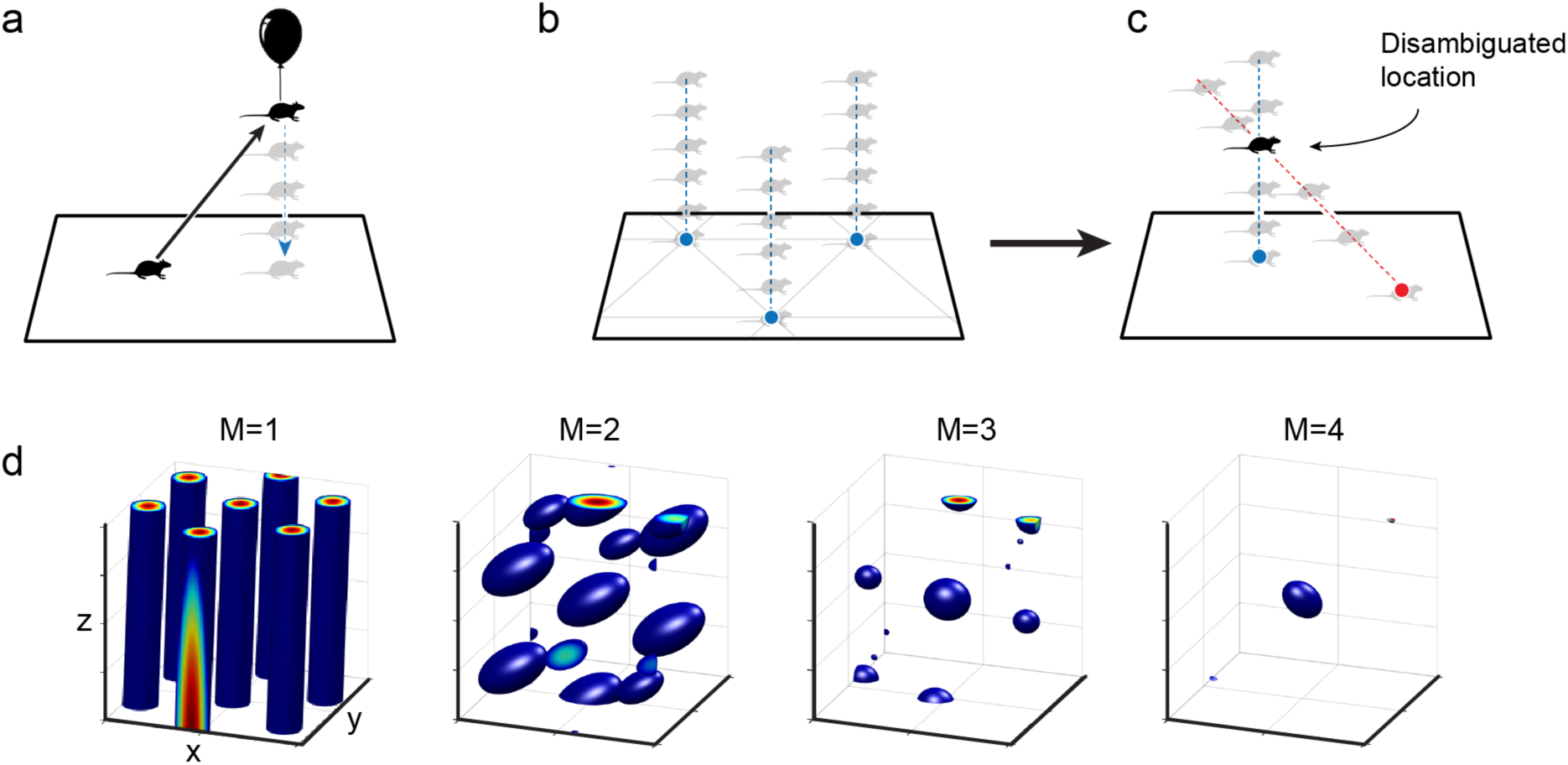
**(a)** If the encoded variable is 3D (here, the animal leaves the 2D plane), simple projection down to the 2D phase is ambiguous and consistent with multiple locations in the z-direction. **(b)** There are two sources of ambiguity, the periodicity of the grid code within in the *xy*-plane, and the ambiguity the z-direction. **(c)** Two different 2D phases for two modules are set by two distinct projections (red and blue) of the 3D value onto a plane. Together they are able to simultaneously resolve both sources of ambiguity. **(d)** Estimates of the value of the encoded 3D variable obtained by combining the ambiguous estimates of 1, 2, 3, and 4 modules as in (c). Given the cell responses we compute a probability estimate and show only areas that exceed a fixed threshold (blue blobs). The spacing between the blobs defines the coding range, that is, the range over which the code is unique. With increasing number of modules the range quickly grows larger than the individual periods.

The central observation of this paper is that just as multiple modules operating independently to integrate velocity solve the problem of the ambiguity of representation by periodic responses, they can *simultaneously* solve the ambiguity that results from the compression of higher-dimensional inputs to two-dimensional responses: We propose that the grid cell system might uniquely encode higher-dimensional variables by constructing *M* distinct operators *A*_*α*_ (of size 2 × *N*) that project velocities in *N*-dimensional spaces into 2-dimensional velocity signals for each of the *M* grid modules. These velocity signals, distributed to different modules, are not related simply by a scalar gain, as for 2D spatial responses, but must differ more fundamentally in their responses to each input dimension (Fig. 2c). We choose *A*_*α*_ to be a random projections independently chosen for each module. These *M* independent projections can be viewed as a single matrix of size 2*M* × *N*, which is of full-rank almost surely if *N* ≤ 2*M* (Appendix). This implies that the intersection of the kernels of all projections is trivial, consisting solely of the zero vector. In consequence they compress different portions of the input space and can mutually resolve their ambiguities (Fig. 2c,d). With realistic estimates for the number of grid modules in an individual animal (*M* = 4*,…*, 8), the code could represent variables that lie in quite high-dimensional spaces (*N* = 8*,…*, 16).

As we show and explain next, it is a remarkable attribute of the grid code that with this simple random projection scheme, the system can flexibly deploy and trade-off its massive coding capacity between coding range per dimension and number of dimensions, across variables of different dimensions.

### 2.4 Expected coding range of the model

In any coding space, the available volume grows exponentially with the number of coding dimensions. We call an *efficient code* one that utilizes a sizable fraction of the available volume for coding. The remarkable property of the grid cell code, enabled by its set of periodic modules, is that it generates a vast library of codewords that are both well-spaced and sufficiently densely packed in the available volume, so that the *range* of inputs represented invertibly by the code scales exponentially with the number of modules [7, 21, 19] (cf. also Equation 5 below). Therefore, the grid code is a highly efficient code for 2D analog variables. We can now ask, with regard to our proposed coding model for generating unique codes for high-dimensional variables, is this efficiency lost?

To address this question, we define the *representational range* of the code to be the maximal side-length of a hypercube of dimension *N* over which no two points are assigned the same grid code. Before we present numerical results on the coding range of our model and its efficiency, we set a rough benchmark by defining a conceptually simpler yet efficient coding scheme for a variable *x* of dimension *N* (with 2 < *N* ≤ 2*M* (cf. also Fig. 4, middle panel).

For simplicity, assume that the number of modules is a multiple of *N*. Then we can divide the modules into *N* distinct groups of *M/N* modules. Let each group encode just one of the *N* coordinates of *x*, so that the problem of encoding a variable of dimension *N* has been decomposed into the task of encoding *N* variables of dimension 1 each. The representational range *L* will then be determined by the minimum coding range of one of the groups, that is

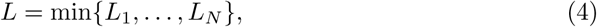

where *L*_*i*_(*i* = 1,…, *N*) denotes the 1-dimensional coding range of the *i*th group.

We know from previous theoretical work [7] that the representational range of encoding a 1-dimensional variable with a number of 2*D* modules grows exponentially in the number of modules. Thus, the representational range *L*_*i*_ of group *i* increases with the number of modules in that group, or as *M/N*. More precisely [7], for a phase resolution (phase bin size). Δ. and an average grid period *>* the expected representational range 𝔼(*L*_*i*_) of group *i* will scale as the spatial size of each phase bin times the number of distinct phases. Thus, we have (cf. Appendix, Figure 6b)

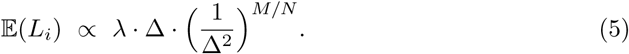

(An expected value is taken because of a random choice of how to “slice” each 2D grid module to represent the *i*th component of *x*.) When *M* is a multiple of *N* we can compute the empirical distribution for *L*_*i*_ (Appendix, Figure 6a) as well as an expected value for the overall representational range *L* (Fig. 3, solid lines). For non-integer values *M/N* ≥ 1, we simply interpolate between the distributions of *L*_*i*_.

**Figure 3:**
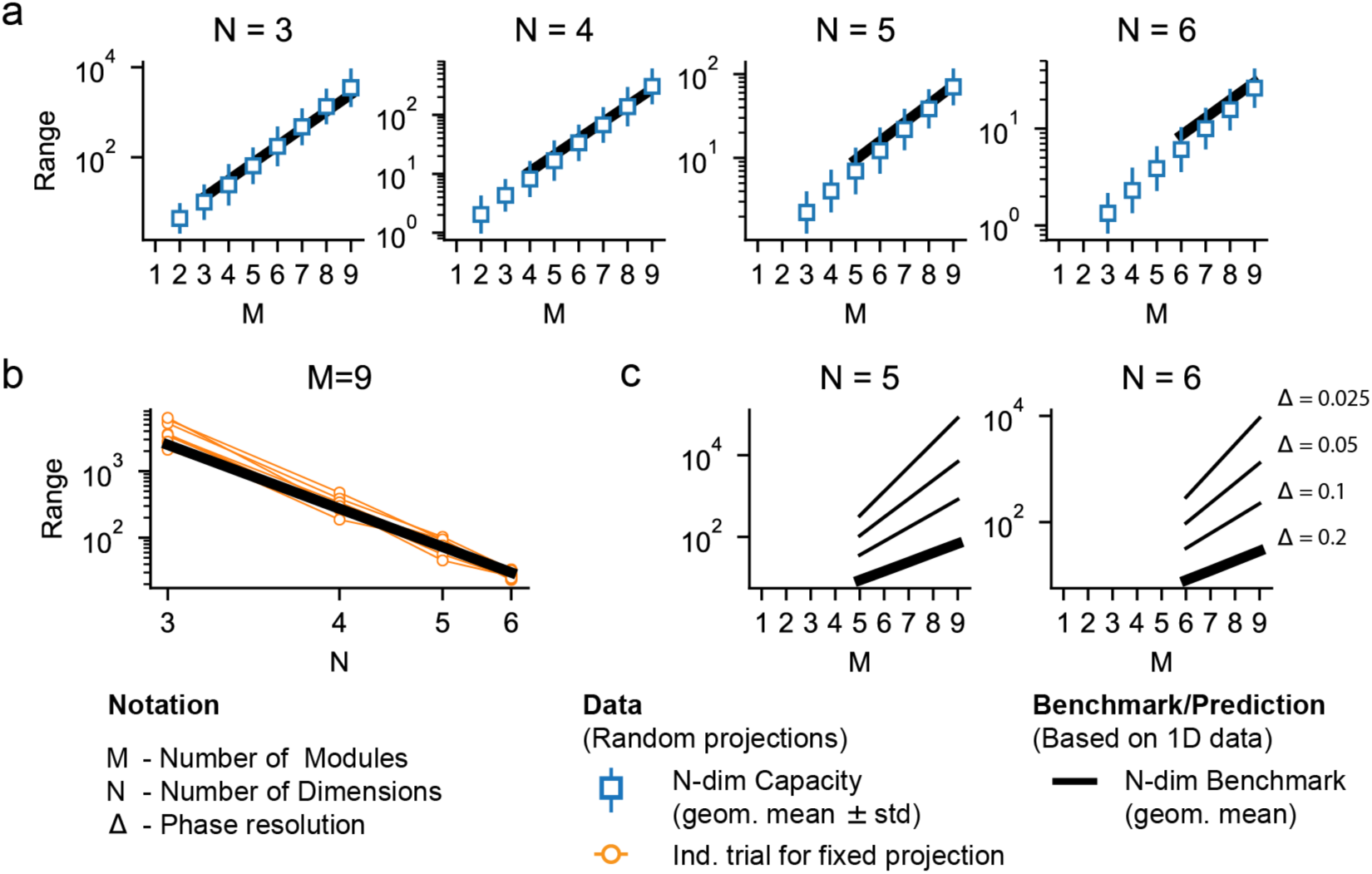
Capacity grows exponentially with module number. All plots show the *dynamic* coding range of our model (see Methods section). **(a)** Exact coding range of the grid code for variables of dimension 3 to 6, assuming an overly conservative phase resolution of Δ = 0.2 to reduce time of computation. We show the geometric mean and standard deviation over 1000 different draws of the projection matrices *A* for each pair *M, N*. The entries of the matrices are sampled independently from a standard normal distribution. To compute the expected value E(*W*) of the benchmark in Equation 4, we also run this simulation with *N* = 1 (Appendix, Fig. 6), solid line. The capacity grows exponentially with the number of modules; the benchmark provides an estimate of the expected capacity. **(b)** Change of coding range while increasing the dimensionality of the input and keeeping the projection fixed (per trial), illustrating the flexibility of the scheme (10 trials are shown). **(c)** We use the benchmark to show the coding range for more realistic values of phase resolution (Δ = 0.2,*…*, 0.025). We chose the benchmark rather than measuring the exact range for practical reasons (the run-time scales with the *volume* of the coding range not its side-length). Results shown for *M/N* ≥ 1.

The benchmark shows that the representational range of our model should still increase exponentially with the number of modules when representing higher-dimensional variables. While the benchmark scheme is simple and efficient, it is based on the specific dimension of the input (this determines how modules are grouped) and hence is not *flexible*, in contrast to our model, as we will consider below.

### 2.5 Efficiency and flexibility of the model

For input variables of various dimensions (*N* = 3,*…*, 6) we numerically compute the representational range as a function of the number of encoding grid modules (*M* = 1,*…*, 9), Figure 3a (white squares show the mean value over different samples of the projection matrices). For each module *α* we generate a random set of projections *A*_*α*_ of dimension 2 × 6; these projection matrices remain unchanged as the input dimension *N* is varied (though only the corresponding sub-matrix of dimension 2 × *N* gets used for coding).

The representational range of the randomized scheme clearly grows exponentially with the number of modules, regardless of variable dimension. Further, the rate of exponential growth closely matches the benchmark, illustrating its efficiency. Thus, the 2D grid code with multiple modules is capable not only of unambiguous representation of arbitrary 1D and 2D variables with exponentially large representational range as a function of number of modules, but it can do the same for much higher-dimensional variables as well.

Next, we note the model’s flexibility. When we fix the random projections *A*_*α*_, then increase the dimensionality of the input variable, the same projection enables representation of all additional dimensions of the input as well: The additional dimensions are encoded by appropriating states from the far end of the range of the smaller-dimensional representations, as can be seen because the representational range shrinks (while remaining exponential) as the input dimension is increased, Figure 3b. The conceptual reason behind the inflexibility of the benchmark coding scheme using any modular code including grid cells, versus the flexibility of our random projection-based scheme with grid cells is illustrated schematically in Figure 4 (right panel).

**Figure 4:**
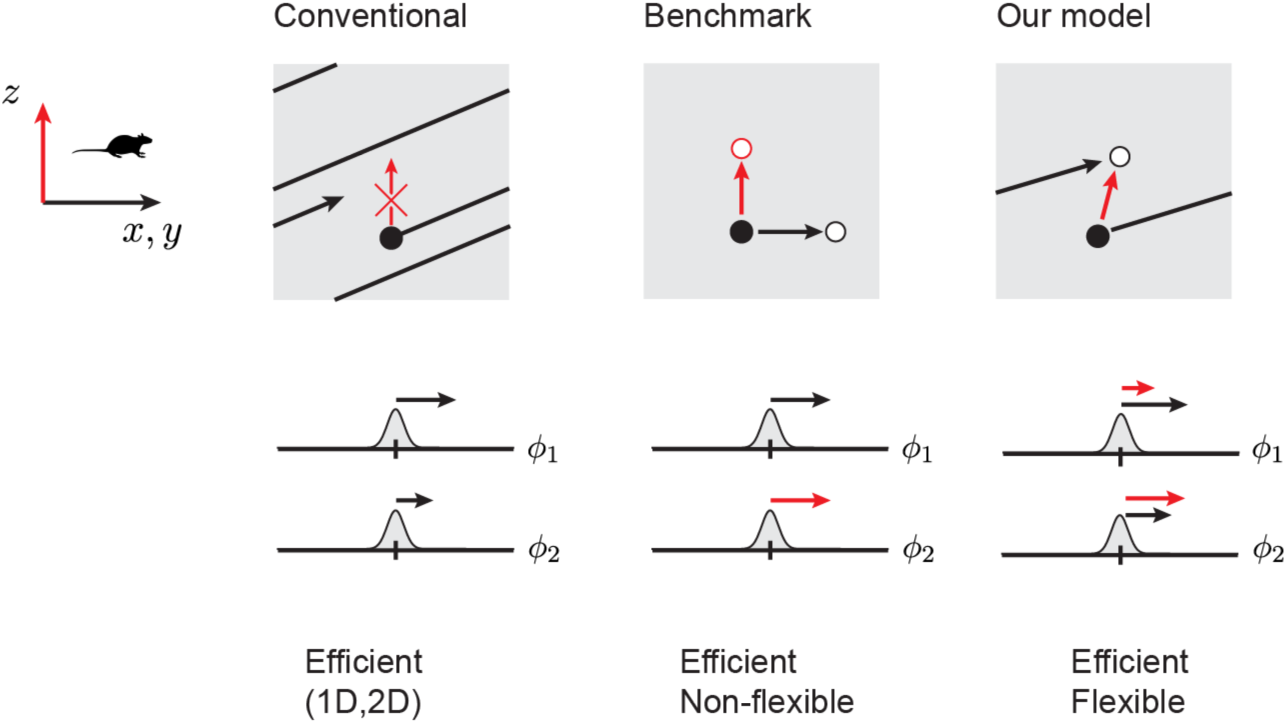
Conceptual view of efficient coding schemes that are flexible versus inflexible. **Left:** The conventional (2D-only) grid code; cf. Figure 1c. Different modules change their state (phase) in a way that differs by a fixed scalar gain; this rigid coupling prevents the positional code from leaving a predefined set of coding states, defined by planes that are arranged in parallel (black lines are edge-on views of these planes) within the combined 2*M*-dimensional phase coding space of the grid cell system. Because the planes are tightly coiled (folded) throughout the coding space (gray), the range is exponentially large in number of modules. However, it cannot encode variables of dimension greater than 2. **Middle:** If modules are partitioned into disjoint groups, and different inputs (*x, y* versus *z*) control the state updates in the different groups, the system can encode higher-dimensional variables. However, the partitioning of modules must be carried out in advance for the specific dimension of the variable: A state that can be reached via updates in one set of inputs (the black direction) cannot be accessed when another set of inputs is varied (red direction), which means each state is pre-allocated to represent a given input dimension. States cannot be traded to exchange coding range for coding dimension, thus such codes are not flexible. **Right:** As in the middle example, updates in module activity are decoupled, but this time *each* module participates in the representation of *all* input dimensions. The periodicity of the code makes it possible to connect (thus use) two coding states in different ways: by moving along different input dimensions (red and black arrow). This means a given state (white circle) can be used to *either* enlarge the coding range in the black direction *or* to encode a new dimension of the variable (via the red direction), without reconfiguration.

For reasons of computational complexity, our primary numerical calculations are performed with a rather conservative phase resolution (Δ = 0.2). To gain a more realistic picture of model performance with finer phase resolution, we consider how coding range of our model changes with phase resolution for a moderate number of modules and dimensions (Appendix, Figure 7) and also how the benchmark changes with phase resolution (Figure 3c). Consistent with [7], capacity grows as a power of phase resolution, regardless of the dimensionality of the encoded variable.

### 2.6 Predicted tuning curves for *N*-dimensional representations

Will it be possible to identify whether grid cells could be performing a flexible representation of high-dimensional variables? A prerequisite for the grid cell system to work according to our model is that different modules must be capable of changing their internal state independently of each other (through the action of separate velocity projection operators); tantalizing hints that this is possible appear in [22], where different grid modules appear to rescale by different amounts in response to an environmental deformation.

A key signature of our proposed scheme relates to differences in predicted tuning curves across modules, described next. While it is straightforward to state the predicted properties of neural responses to *N*-dimensional variables, it is difficult to record, visualize, and characterize these tuning curves in both principle and practice. Fortunately, doing so is not necessary. We can simply characterize the responses of cells in different modules as a function of variations along any 2D subspace of the explored *N*-dimensional input space. The tuning curves in *N* dimensions will look like “lifts” (unchanging responses) of these 2D slices along *N* − 2 of the remaining dimensions (the null space of the projection). For instance, if the input variable is 3D, these lifts simply look like elongated bands or pillars (see Figure 5a, *M* = 1).

**Figure 5:**
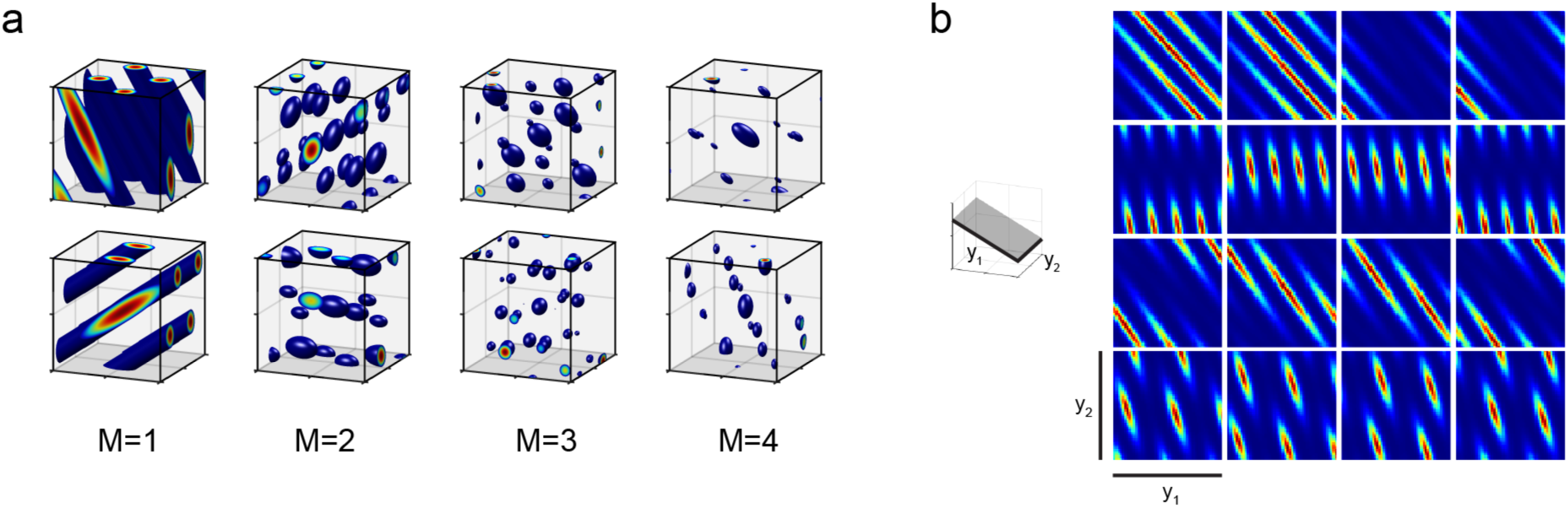
Predictions about grid cell firing. For ease of illustration, we consider here the encoding of a variable in three dimensions. **(a)** Left-most column (*M* = 1): 3D tuning curves of two grid-cells from different modules using our coding model. Remaining columns (*M >* 1): 3D tuning curves of two conjunctive cells reading from *M* different modules using our coding model. **(b)** Each row shows the 2D responses of 4 co-modular cells over a randomly chosen tilted plane (shown on the left in gray) in 3D space. Different rows correspond to different modules and the modules encode space according to our model.

The responses of grid cells along the 2D subspace will then resemble distorted grids (Figure 5b), and can range from perfectly equilateral triangular grids to non-grid-like and relatively complex, for instance, bands of bands (Figure 5b, 2nd row); in the atypical case where the 2D subspace exactly aligns with one of the null or lift directions of a module, those cells will have periodically arranged stripe responses. For a broader sampling of possible grid cell response geometries, see Appendix Figure 8. Within a module, the responses of different cells are generated from translations of the tuning of the module (each row of Figure 5b shows co-modular responses over 2D slices). If plotted over a large enough area, these translational relationships will be apparent, but when plotted over smaller areas, they need not appear as simple shifts of a canonical 2D response pattern (e.g. Figure 5b, top row), similar to the relationships seen in co-modular cells in 1D environments which are generated by cutting a lower-dimensional slice through translations of a higher-dimensional (2D) lattice [29]. Nevertheless, the common origins of the response of co-modular cells means that they will obey systematic relationships.

Many cells in entorhinal cortex and the hippocampus express spatial fields that are less structured than grid cells. In our model, if some of these cells were performing a readout of two or three modules, these conjunction-forming cells would exhibit localized 3D fields with some regularity in spacing, but not full grid-like periodicity and thus no clear notion of a spatial phase (Fig. 5a).

In sum, a central prediction of the hypothesis that the grid cell system could collectively and flexibly use its multiple modules to encode variables of higher dimension than two is that the projections to different modules should be different, and therefore that in such situations, the responses of grid cells in different modules will differ in the geometry of their tuning curves.

## 3 Discussion

### 3.1 Implications for computation

The multi-module representation of grid cells provides a pre-fabricated, ready-to-use, general high-dimensional neural affine^1^ vector space that can be used for both representation and memory of arbitrary vectors (of dimension ≤ 2*M*), and more specifically, for integration of vector inputs. The representation is efficient: it generates exponentially many states with linearly many neurons, thus solving the curse of dimensionality problem faced by more naive coding schemes (e.g. by the formation of unary codes or grids in higher dimensions). The update mechanism of grid cells permits vector-algebraic operations between the stored vectors, required for vector integration in higher-dimension abstract spaces. So long as displacements in the abstract spaces are provided as inputs to the network, the network can thus efficiently represent, hold in memory, and perform algebraic sum operations on general, abstract vectors of different dimensions without any reconfiguration of the recurrent grid cell network. We believe these results and implications fulfill, at least in theory, intuitive expectations that the very peculiar grid code might be extraordinary in the computations it enables.

### 3.2 Relationship with band cells

Cells whose spatial responses resemble evenly spaced parallel bands across a 2-dimensional environment are called *band cells*. They were postulated as inputs for computational models of grid cells [5], and later observed experimentally in recordings in the parasubiculum and entorhinal cortex of freely moving rats [18]. In our model, tuning curves can be variously grid-like, distorted or stretched grid-like, band-like, amplitude-modulated band-like, or like bands of bands, in a single cell. The band-like responses in our model – obtained from 2D slices through the *N*-dimensional tuning curve – are a generalization of simple band cells, and suggest that previously observed band-like responses might be a signature of a projection operator that projects a 2D or higher-dimensional input onto one of the null directions of the grid module.

### 3.3 Observed 3D responses in grid cells

In some studies of animals exploring higher-dimensional spaces, specifically 3D spatial environments, the response of grid cells is elongated and undifferentiated along one dimension, while remaining grid-like in the other two [11]. This kind of tuning is consistent with our prediction, and we have shown it allows for unique coding along the third dimension if the projections (and thus the undifferentiated direction) are not aligned across modules.

Recently, grid cell responses have been examined in bats flying through 3D environments [9]. Bats crawling on 2D surfaces exhibit the same 2D triangular grid cell tuning [27] as rats and mice. In 3D, consistent with our theory, the responses seem not to clearly exhibit regular 3D grid patterns [26]. However, the fields do seem to be localized in all 3 dimensions, at least in the vicinity of a tree around where the bats forage for food [9]. It is possible in this case that localized higher-dimensional fields are formed in the hippocampus or the lateral entorhinal cortex based on spatial landmarks. Alternatively, localized fields seen in medial entorhinal cortex and hippocampus in 3D could be formed by conjunctions of grid cells encoding higher-dimensional spaces according to our model, as shown in Fig. 5a, which qualitatively matches some of the reported properties of entorhinal cells in flying bats. A similar situation might hold for the observed localization of fields in 3D, in rats navigating 3D wire mesh cubes [14]. However, an absence of band-like structure in grid cells along any dimension during 3D coding would not be consistent with our theory.

## 4 Methods

For the numerical part of the paper, we constructed a different random projection for each trial. The entries of the matrices were independently sampled from a standard normal distribution. Given a collection of *M* grid modules and their associated projection matrices, we want to determine the maximal range over which the code is unique. We say that two points *x, x′ collide* if the distance of their associated phases in the grid representation is smaller than or equal to a fixed threshold 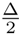, where Δ is the phase resolution per module. We determine the capacity of the code by computing the side length of a maximal collision-free cube centered at the origin. However, in a small neighbourhood of the origin (moving along each dimension by an amount smaller than all the grid periods) the encoding map is one-to-one and continuous if the intersection of the kernels of the different projection matrices is trivial. Any point thus admits a small neighbourhood of points whose associated phases are closer than Δ; it is necessary to ignore these points while performing our search for collisions for the capacity computations. We compute the minimal hyper-rectangle enclosing the ignored points and then incrementally extend this box outward. We define the *dynamic range* as the minimum of the side-lengths of the maximal cube divided by the side-length of the minimal starting box. See Supplementary Material for a more detailed description of the algorithm.

## Acknowledgements

This work was initiated at the Simons Institute for the Theory of Computing at the University of California, Berkeley. IRF is an HHMI Faculty Scholar and was funded in part by the Simons Foundation through the SCGB Collaboration.

## Footnotes

1 The term *affine* makes explicit the lack of a preferred “zero” element. Each point in the space admits a neighbourhood that naturally carries the structure of a vector space with the point at its origin.

