## Appendix for "Flexible representation of higher-dimensional cognitive variables with grid cells"

### Appendices

#### A Methods

#### B Rank of random projection is maximal

In the paper we state that the random projection $A \in \mathbb{R}^{2M \times N}$ has rank $N$ with "high" probability if $N \leq 2M$. We will show below that the set of $(2M \times N)$-matrices of rank less than $N$ can be expressed as the zero set of a polynomial (which is not the zero polynomial). Thus its measure is zero and hence the probability of sampling one of these lower-rank matrices is zero as well. One way to construct such a polynomial is as follows. For $I = (i_1, \ldots, i_N)$ let $A_I$ denote the $(N \times N)$-matrix formed by the rows of $A$ whose indices are listed in $I$, i.e. the entry in row $k$ and column $l$ of $A_I$ is given by $a_{i_k,l}$. Recall that for any square matrix we can compute its determinant, and recall further that the determinant is a polynomial expression. With this in hand we can now define a polynomial function $f$ on the set of $(2M \times N)$-matrices whose zero set is given by matrices of rank less than $N$:

$$f(A) := \sum_{1 \leq i_1 < \ldots < i_N \leq N} \det(A_I)^2.$$

Note that $f(A)$ is non-zero, if and only if at least one of the summands is non-zero. The latter is equivalent to $N$ of the $2M$ rows of $A$ being linearly independent, which means that the rank is $N$.

#### C Existence of preferred directions

For any projection matrix $A \in \mathbb{R}^{2 \times N}$ of maximum rank $N$ there exist two *preferred* non-null directions such that the responses look perfectly triangular. This translates to the problem of finding a pair of vectors $v, w \in \mathbb{R}^N$ that satisfies the following properties: (i) They map to the standard basis in $\mathbb{R}^2$, i.e. $A(v) = e_1$ and $A(w) = e_2$; (ii) They are orthogonal, i.e. $\langle v, w \rangle = 0$; (iii) Their length is equal, i.e. $\|v\|^2 - \|w\|^2 = 0$. We can already see that we can expect such a pair to exist. Note that all three properties amount to 6 equations and our hypothesis space is $2N$-dimensional since it is the product of two $N$-dimensional Euclidean spaces. Thus for $N > 2$ we have just enough degrees of freedom to expect a solution to exist. For $N = 3$ let $V$ and $W$ denote the solutions to the equations in (i). For $N > 3$ we can choose 1-dimensional, parallel, affine-linear sub-spaces of the solutions to equation in (i). For $v \in V$ let $w(v)$ denote the intersection of the orthogonal complement of $v$ and $W$. This map is well-defined if $v$ is not orthogonal to $\ker A$. So far the first two of the three properties are already satisfied. If we let $\|v\|$ go to infinity, $\|w(v)\|$ will approach a minimum, and in turn approaching the the orthogonal complement of $\ker A$ with $v$ will let $\|w(v)\|$ approach infinity. Thus following a mean value theorem argument one concludes that there is a $v$ with $\|v\| = \|w(v)\|$.

#### D Details of the capacity algorithm

We determine the side length $w$ by first determining the *resolution* of the set of modules (discussed later) and confirming that it is less than 0.5 units in each dimension. We then started with a small $N$-dimensional box of radius 0.5, and we incrementally expand the box outward and check whether the frontier region contains any collisions, splitting this frontier into a set of $N$-dimensional boxes and thoroughly checking each of these boxes for the starting point's representation. There's no analytic way to detect whether a particular set of module phases occur together within a $N$-dimensional box, but in some cases it's trivial. For example, if one of the phases never occurs anywhere in the box (while accounting for $\Delta$), then it's clear that the set of phases never co-occur in the box. We combine this heuristic with a divide-and-conquer algorithm to split the box into smaller boxes in which at least one phase never occurs. For each box we test a few points in the box to see if they have this representation, then we test whether the box excludes any of the individual phases, and if both checks fail we split the box in half and try again on each half. If the representation is not present, this process will successfully partition the box into smaller boxes that each have this property. If the representation does occur, this process will narrow in on a range of points with representations similar to this representation until it finds it. In this way, we thoroughly search the N-dimensional volume for collisions without needing to choose a fixed high sampling density.

In each module we anchor $\phi_0 = \vec{0}$ to location $\vec{x}_0 = \vec{0}$, so $A$ assigns phase $\vec{\phi}$ to a location $\vec{x}$ via $A\vec{x} \mod 1$. In the algorithm above we need to check whether various $N$-dimensional boxes contain any points near phase $\vec{0}$. To perform this check in a way that properly considers distances between phases, we decompose $A$ into two matrices: $P$, a linear mapping from $\mathbb{R}^N$ to the plane $\mathbb{R}^2$, and $L$ which specifies the lattice on that plane, such that $A = L^{-1}P$. We always set $L$ to create a hexagonal lattice, i.e.

$$L = \begin{bmatrix} \cos 0° & \cos 60° \\ \sin 0° & \sin 60° \end{bmatrix}. \tag{6}$$

The matrix $P$ maps a location to a point on the plane that contains the hexagonal lattice:

$$P = \frac{1}{\lambda} \begin{bmatrix} 1 & 0 & 0 & \cdots & 0 \\ 0 & 1 & 0 & \cdots & 0 \end{bmatrix} \begin{bmatrix} \hat{v}_1 & \hat{v}_2 & \cdots & \hat{v}_N \end{bmatrix}^{-1}. \tag{7}$$

Here $\hat{v}_1$ and $\hat{v}_2$ are orthogonal $N$-dimensional unit vectors that define this plane, and $\hat{v}_3, \ldots, \hat{v}_N$ define the kernel of $P$. The period or scale of the grid is defined by $\lambda$. The distance between two phases is equal to the shortest distance between the phases on this plane.

To determine whether an $N$-dimensional box contains a point with a phase near $\vec{0}$, we apply transformation $P$ to each corner of the box to obtain its shadow on the plane. We enumerate nearby lattice points on the plane, drawing circles around each lattice point with diameter equal to the phase resolution $\Delta$, and then we check whether any of these circles intersect the shadow or are contained within the shadow. We combine this technique with the above "divide-and-conquer" and "expanding box" techniques to determine the range over which the code is unique.

We defined the *resolution* of a set of modules as the distance one must move along each dimension before the representation becomes distinguishable from the starting representation. It's possible for this resolution to be more precise in some dimensions than others, for example if all of the modules have similar projection planes and similar kernels the resolution on the projection plane will be precise but along the kernel it will be more coarse. To obtain a single number characterizing this resolution, we numerically computed the smallest N-dimensional hypercube centered at the origin for which every point on the hypercube's surface was distinguishable from the origin. We checked each face of the hypercube using the same divide-and-conquer strategy as above. The resolution is equal to half the side-length of this smallest hypercube, and we computed it to a

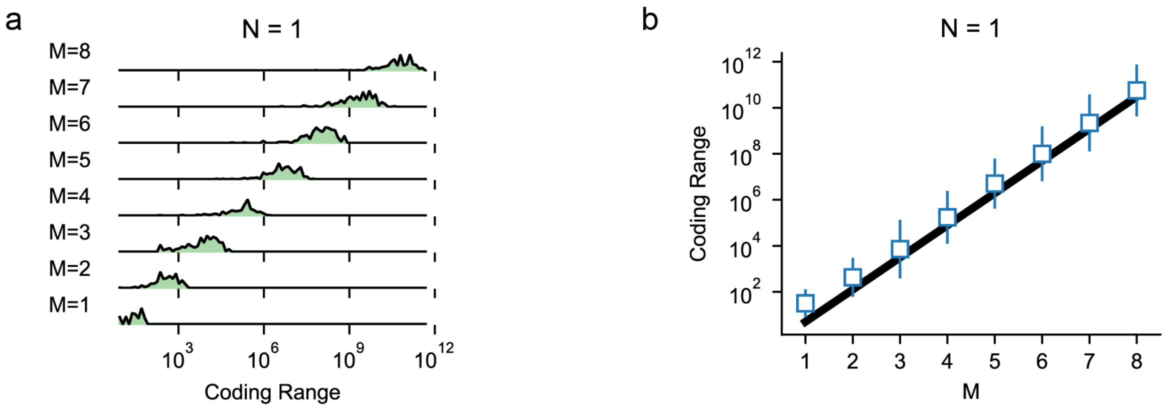


Figure 6: **1-dimensional capacity.** **(a)** Histograms of the 1D capacity data. Note that for each $M$ the distribution is roughly log-normally distributed (1000 data-points for each $M$). For all computations the phase resolution is $\Delta = 0.2$. **(b)** The 1D capacity of our randomized approach (blue error-bars). We show geometric mean and standard deviation of the data (1000 data-points for each $M$). The capacity grows proportional to the benchmark $\Delta \cdot (1/\Delta^2)^M$ (thick black line); cf. [7].

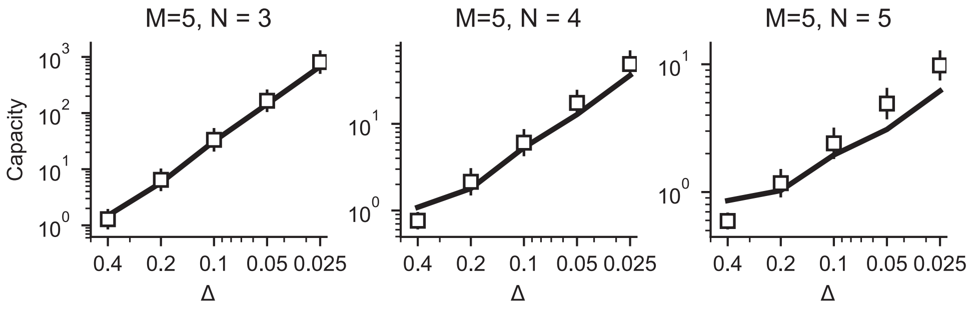


Figure 7: **Capacity as a function of phase resolution.** Capacity grows as a power of phase resolution, regardless of the dimensionality of the encoded variable

precision of 0.01 units.

#### D.1 Phase distance

We base our distance computations in the space of the vector of grid phases with respect to the equivalence relation defined by the lattice (each module encodes a 2D phase on a 2D torus, which we can understand as the quotient of Euclidean space with a hexagonal lattice). We now define a metric on the sets of phases by taking the maximum of their component-wise distances, i.e. for two sets of phases $\boldsymbol{\phi} = (\phi^1, \ldots, \phi^M)$ and $\boldsymbol{\psi} = (\psi^1, \ldots, \psi^M)$ we define

$$d(\boldsymbol{\phi}, \boldsymbol{\psi}) = \max\left\{d_1(\phi^1, \psi^1), \ldots, d_M(\phi^M, \psi^M)\right\},$$

where $d_i$ $(i = 1, \ldots, M)$ denote the metrics inherited from $\mathbb{R}^2$ with respect to each module's underlying lattices.

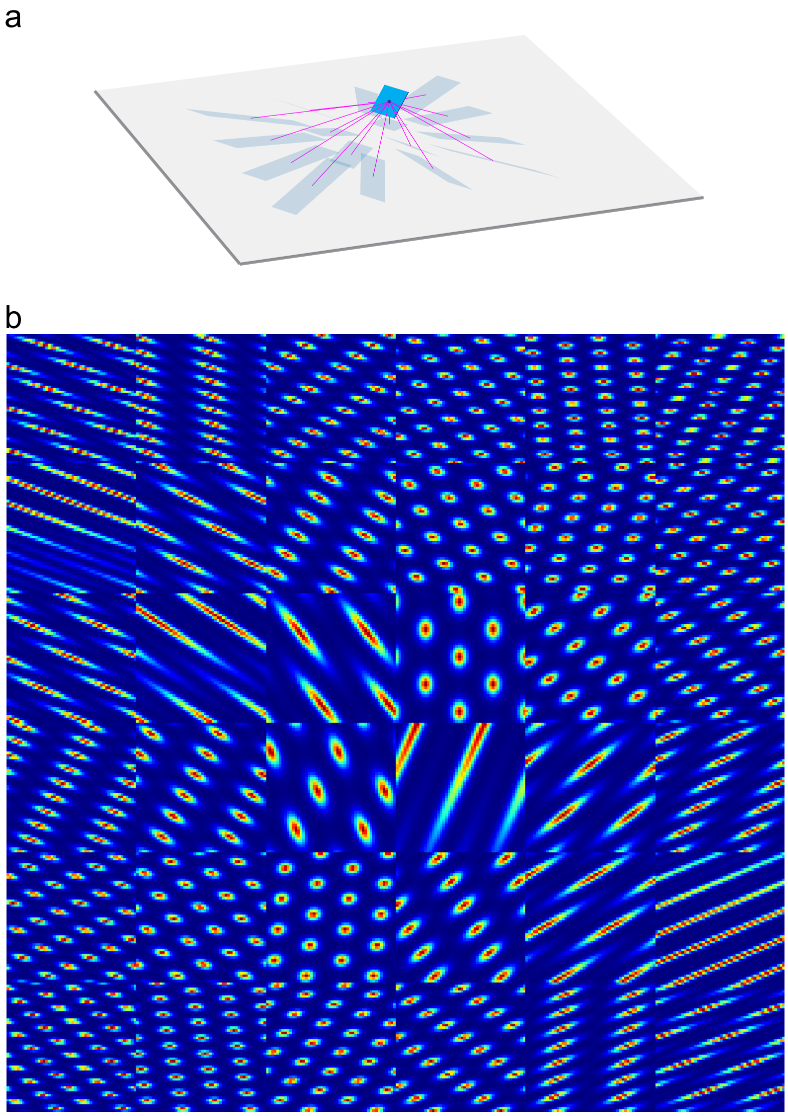


Figure 8: **A zoo of firing fields.** **(a)** A tilted plane (blue) in 3D space and 16 different projections (magenta lines) onto a common 2D input subspace (gray) along which the responses are an equilateral triangular lattice. **(b)** $6 \times 6$ different Firing fields on the blue plane induced by a family of $6 \times 6$ projections whose angles vary as indicated in (a).
